# Genome-Wide Polygenic Risk Scores and Prediction of Gestational Diabetes in South Asian Women

**DOI:** 10.1101/574616

**Authors:** Amel Lamri, Shihong Mao, Dipika Desai, Milan Gupta, Guillaume Paré, Sonia S. Anand

## Abstract

Gestational diabetes Mellitus (GDM) affects 1 in 7 births and is associated with numerous adverse health outcomes for both mother and child. GDM is suspected to share a large common genetic background with type 2 diabetes (T2D). The first aim of this study, was to characterize different GDM polygenic risk scores (PRSs) using data from the South Asian Birth Cohort (START). The second aim of this study was to estimate the heritability of GDM.

PRSs were derived for 832 South Asian women from START using the pruning and thresholding (P+T), LDpred, and GraBLD methods. Weights were derived from multi-ethnic (Mahajan *et al*., 2014) and white Caucasian (Scott *et al*., 2017) studies of the DIAGRAM consortium. Association with GDM was tested using logistic regression. Heritability of GDM was estimated using the GRMEL approach. Results were replicated in samples from the UK Biobank (UKB) study.

The top P+T, LDpred and GraBLD PRSs were all based on Mahajan *et al*. The best PRS was highly associated with GDM in START (AUC=0.62, OR=1.60 [95% CI=1.44–1.69]), and in South Asian (AUC=0.65) and white British (AUC=0.58) women from UKB. Heritability of GDM approximated 0.55±0.83 in START and 0.18±0.22 in white British women from UKB.

Our results highlight the importance of combining genome-wide genotypes and summary statistics from large multi-ethnic studies to optimize PRSs in South Asians.

## INTRODUCTION

Gestational diabetes mellitus (GDM) is defined as dysglycemia due to elevated blood glucose levels first identified during pregnancy, and is specifically defined based on glucose response to an oral glucose challenge test in pregnancy. GDM has been associated with numerous adverse health outcomes affecting mother and child, both during and after pregnancy ^1,2^. Because of its increasing prevalence (∼ 1 in 7 births), GDM has become a major health concern worldwide ^3^. Nevertheless, the prevalence of GDM largely varies from one region of the globe to the other, and South Asian women have been shown to be at higher risk of GDM than white British women.

Although GDM is thought to have a strong genetic component, to our knowledge, no studies have estimated the heritability of GDM. Nevertheless, since GDM and T2D have similar risk factors and share common pathophysiological pathways, the heritability of GDM is thought to be similar to that of T2D.

Numerous genome-wide association studies (GWASs) and genome-wide association meta-analysis (GWAMAs) of glucose related traits and T2D have been conducted in non-gravid populations, and summary statistics from large consortia (e.g., MAGIC and DIAGRAM) are publicly available ^4-13^. By contrast, few studies of genetic determinants of GDM have been conducted or published. For instance, only two studies sought to identify genes associated with dysglycemia, GDM, and diabetes during pregnancy by GWAS ^14,15^. Top signals from these studies were located within/near *CDKAL1, MTNR1B, GCKR, PCSK1, PPP1R3B* and *G6PC2*, which were previously known for their association with glucose metabolism and T2D ^14,15^. In addition, other T2D associated loci (e.g., *TCF7L2, PPARG, CDKN2A/B, KCNQ1, GCK*, etc.) were also significantly associated with GDM when tested separately ^16-39^, or combined in genetic risk scores (GRS) ^33,34,40-42^.

GRS are used to capture genetic information at one or more loci. Most of published studies interested in complex traits/diseases and using GRS typically combine data for a small number of single nucleotide polymorphisms (SNPs), and the predictive power of these GRS is sub-optimal ^43^. However, with the increased availability of genome-wide genotypes and publicly available data from large consortia, GRS with a larger number of variants are being used, and the predictive value of these genome-wide polygenic risk scores (PRSs) has substantially improved ^44,45^.

PRSs can be derived using different approaches, however, these require both summary statistics from an external GWAS and genetic data for a reference panel for between-variants linkage disequilibrium LD (LD) calculations. Pruning and thresholding (P+T) is a commonly used heuristic approach to derive PRSs in which variants are filtered based on an empirically determined P-value threshold. Linked variants are further clustered in different groups and SNPs with the highest significance (lowest P values) in each group are prioritized and included in the PRS, while variants of less significance within the group are pruned out ^46^. Other programs have been shown to improve the predictive value of the score by allowing the inclusion of a larger number of independent as well as linked variants into the score using different approaches. For instance, LDpred, another commonly used method, estimates the mean weight of each variant, assuming a prior knowledge of the genetic architecture of the trait (fraction causal), and using a Bayesian approach ^47^. More recently, we developed the gradient boosted and LD adjusted (GraBLD), a new PRS building approach which applies principles of machine-learning to estimate SNP weights (gradient boosted regression trees), and regional LD adjustment ^48^.

The first objective of this study was to determine the optimal gene scores and investigate the association of genetic variants combined in these PRSs with GDM in South Asian women participating to the South Asian Birth Cohort (START). We considered several parameters: 1) two different sources of summary statistics, namely Mahajan *et al*., 2014 (trans-ethnic GWAMA) ^5^ *vs*. Scott *et al*., 2017 ^6^ (GWAMA in white Caucasians); 2) two templates for LD calculation (1000 Genomes phase 3 ^49^ *vs*. START genotypes); 3) different minimal values of the number of samples in each SNP’s analysis in the consortia; 4) weighted *vs*. unweighted PRSs; 5) three methods to derive the PRSs; Pruning and Thresholding (P+T),. LDpred and GraBLD and; 6) different P-value thresholds (for P+T and LDpred PRSs only). The second objective was to estimate the heritability of GDM from: 1) genome-wide data; 2) common variants; and 3) SNPs in the best P+T PRS. PRS results were further validated in South Asian women from UK Biobank, and both PRS and heritability results were replicated in a subset of white British women from UK Biobank^50^.

## RESULTS

### Population characteristics

Table 1 shows the characteristics of START and UK Biobank women included in the analyses. Because of major differences in recruitment strategies, inclusion criteria and study protocols, selected participants from the UK Biobank were of older age, higher weight, and body mass index (BMI) compared to START participants. Furthermore, the proportion of participants with GDM was significantly lower in South Asian women from the UK Biobank and even more so in European women of the same study.

**Table 1:**
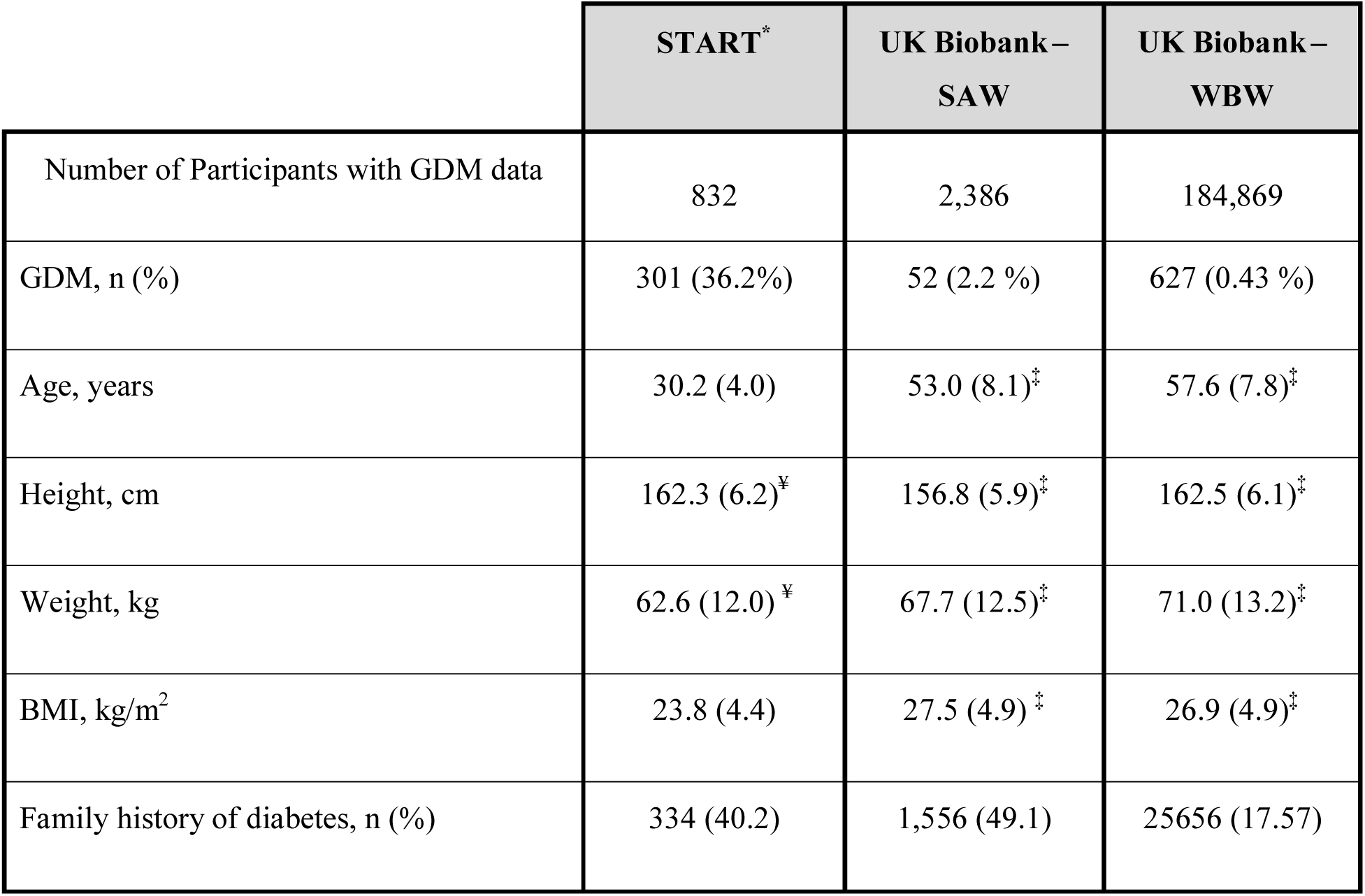
Characteristics of women participants from the START and UK Biobank studies with available GDM and Genotype data. ^*^South Asian Women with GDM status. Data are mean (SD) unless otherwise indicated. ¥ pre-pregnancy values, ^‡^ Values from baseline data. Abbreviations: BMI, Body mass index; GDM, gestational diabetes; SAW, South Asian women, WBW, white British women; START, South Asian Birth Cohort.

|  | START* | UK Biobank –<br>SAW | UK Biobank –<br>WBW |
| --- | --- | --- | --- |
| Number of Participants with GDM data | 832 | 2,386 | 184,869 |
| GDM, n (%) | 301 (36.2%) | 52 (2.2 %) | 627 (0.43 %) |
| Age, years | 30.2 (4.0) | 53.0 (8.1)<sup>‡</sup> | 57.6 (7.8)<sup>‡</sup> |
| Height, cm | 162.3 (6.2)<sup>¥</sup> | 156.8 (5.9)<sup>‡</sup> | 162.5 (6.1)<sup>‡</sup> |
| Weight, kg | 62.6 (12.0) <sup>¥</sup> | 67.7 (12.5)<sup>‡</sup> | 71.0 (13.2)<sup>‡</sup> |
| BMI, kg/m<sup>2</sup> | 23.8 (4.4) | 27.5 (4.9) <sup>‡</sup> | 26.9 (4.9)<sup>‡</sup> |
| Family history of diabetes, n (%) | 334 (40.2) | 1,556 (49.1) | 25656 (17.57) |

### Minimum sample size per SNP in consortium data

Summary statistics were derived from a trans-ethnic (Mahajan *et al*., 2014 ^5^) and white Caucasian (Scott *et al*., 2017 ^6^) GWAMAs of the DIAGRAM consortium. In each study, the number of participants tested for association with T2D was different for each SNP, and ranged between 25 – 110,219 and 4,731 – 158,186 in Mahajan *et al*. and Scott *et al* respectively (Supplementary Figure 3, Supplementary Table 1).

We derived several PRSs for which the list of variants was restricted to SNPs tested in at least 0, 85, 90, 95 and 98% of the maximum sample size in the consortium GWAMA of interest. The number of SNPs used in the PRSs and the percentage of SNP loss for each one of these thresholds are shown in Supplementary Table 1. The percentage of variants lost after this filtering was the most substantial in PRSs based on Mahajan *et al*., with only 346,290 polymorphisms remaining when keeping variants tested in ≥ 95% samples (Supplementary Table 1).

For both Mahajan *et al*. and Scott *et al*. based PRSs, the optimal minimum percent of participants to keep varied depending on the method used (P+T, LDpred or GraBLD), the source of LD estimates (LD_START_ or LD_1KG_) and the consortium P-value threshold (Figure 1, Figure 2, Supplementary Figure 3 and Table 2).

**Table 2:** Characteristics and GDM association results of the top weighted P+T and GraBLD PRSs in South Asian women from the START and UK Biobank studies. Results are from univariate association tests with GDM. The top 3 PRSs are shown in bold. Abbreviations: 1KG, 1000Genomes; AUC area under the curve; GraBLD, Gradient Boosted and LD adjusted; LD, Linkage Disequilibrium; NA, Non applicable; P+T, pruning and thresholding; PRS, Polygenic Risk Score; SAW, South Asian Women; SE, Standard Error; START, South Asian Birth Cohort; WBW, White British Women.

|  |  |  | START - SAW |  |  |  | UK Biobank - SAW |  |  |  | UK Biobank - WBW |  |  |  |
| --- | --- | --- | --- | --- | --- | --- | --- | --- | --- | --- | --- | --- | --- | --- |
| PRS Type | Consortium | LD source | Beta | SE | P-value | AUC | Beta | SE | P-value | AUC | Beta | SE | P-value | AUC |
| P+T | **Mahajan et al, 2014** | **1KG** | **0.445** | **0.08** | **$8.7 \times 10^{-9}$** | **0.62** | **0.423** | **0.14** | **0.003** | **0.61** | **0.303** | **0.04** | **$4.63 \times 10^{-14}$** | **0.58** |
| | | START | 0.448 | 0.08 | $6.55 \times 10^{-9}$ | 0.62 | 0.512 | 0.14 | 0.0003 | 0.64 | 0.291 | 0.04 | $3.84 \times 10^{-13}$ | 0.58 |
| | Scott et al, 2017 | 1KG | 0.370 | 0.07 | $7.86 \times 10^{-7}$ | 0.60 | 0.280 | 0.14 | 0.05 | 0.57 | 0.296 | 0.04 | $1.45 \times 10^{-13}$ | 0.58 |
| | | START | 0.351 | 0.07 | $2.98 \times 10^{-6}$ | 0.60 | 0.300 | 0.14 | 0.03 | 0.59 | 0.270 | 0.04 | $1.88 \times 10^{-11}$ | 0.57 |
| GraBLD | **Mahajan et al, 2014** | 1KG | 0.465 | 0.08 | $1.8 \times 10^{-9}$ | 0.62 | 0.520 | 0.14 | 0.0003 | 0.64 | 0.343 | 0.04 | $1.14 \times 10^{-17}$ | 0.59 |
|  |  | **START** | **0.470** | **0.08** | **$1.32 \times 10^{-9}$** | **0.62** | **0.510** | **0.14** | **0.0005** | **0.63** | **0.351** | **0.04** | **$1.86 \times 10^{-18}$** | **0.59** |
| | Scott et al, 2017 | 1KG | 0.317 | 0.07 | $1.61 \times 10^{-5}$ | 0.59 | 0.388 | 0.14 | 0.006 | 0.61 | 0.344 | 0.04 | $8.08 \times 10^{-18}$ | 0.59 |
| | | START | 0.342 | 0.07 | $3.93 \times 10^{-6}$ | 0.59 | 0.387 | 0.14 | 0.006 | 0.61 | 0.348 | 0.04 | $4.15 \times 10^{-18}$ | 0.59 |
| LDpred | **Mahajan et al, 2014** | **1KG** | **0.461** | **0.07** | **$2.18 \times 10^{-9}$** | **0.62** | **0.527** | **0.14** | **0.0002** | **0.65** | **0.305** | **0.04** | **$2.63 \times 10^{-14}$** | **0.58** |
| | | START | 0.470 | 0.08 | $1.32 \times 10^{-9}$ | 0.62 | 0.44 | 0.14 | 0.002 | 0.61 | 0.278 | 0.04 | $3.32 \times 10^{-2}$ | 0.57 |
| | Scott et al, 2017 | 1KG | 0.347 | 0.07 | $4.05 \times 10^{-6}$ | 0.59 | 0.382 | 0.14 | 0.006 | 0.61 | 0.377 | 0.04 | $1.80 \times 10^{-21}$ | 0.60 |
| | | START | 0.281 | 0.07 | 0.00015 | 0.57 | 0.225 | 0.13 | 0.10 | 0.55 | 0.323 | 0.04 | $1.90 \times 10^{-16}$ | 0.59 |

**Figure 1:**
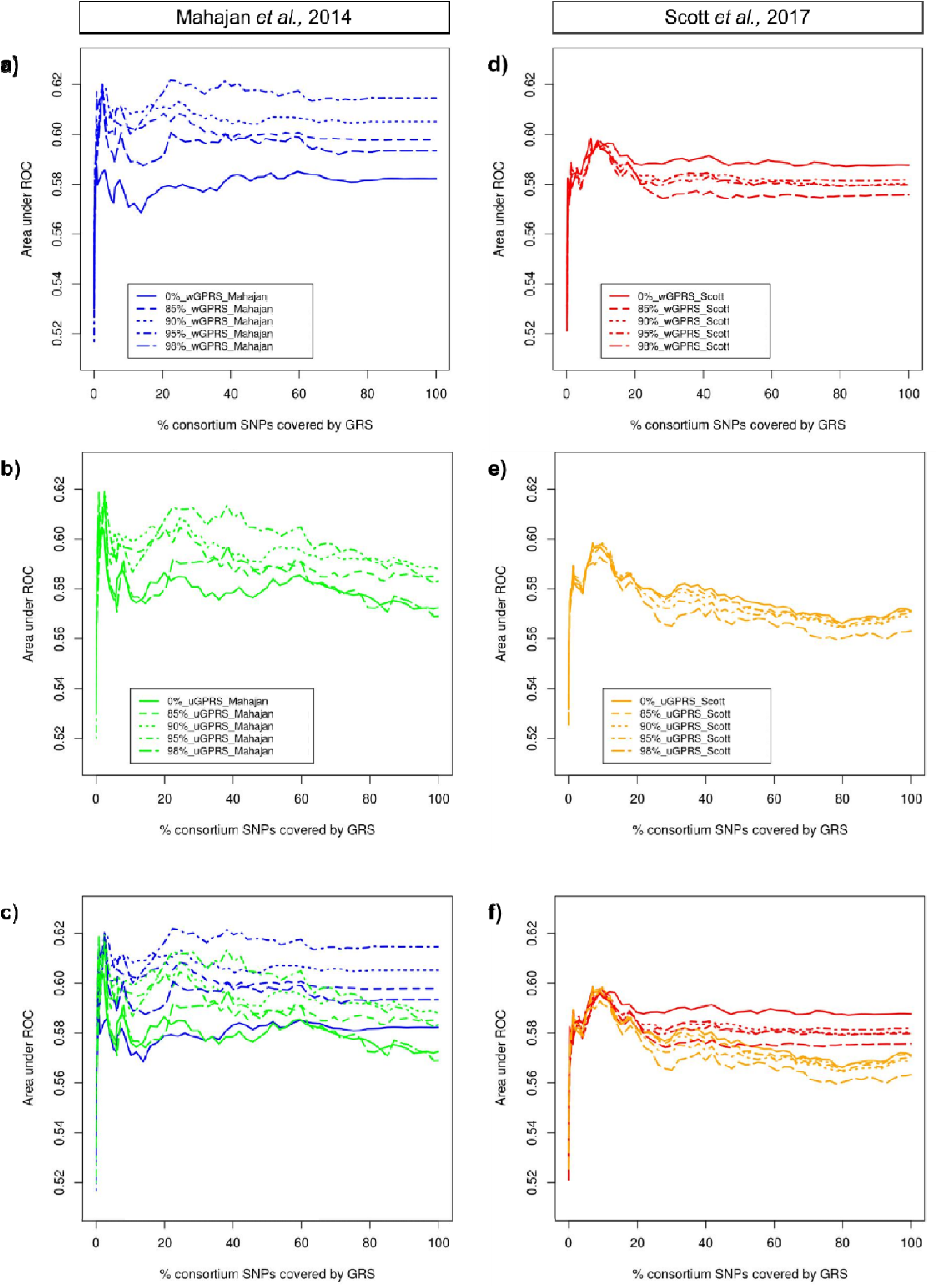
AUCs of the different weighted and unweighted LD_START_ P+T PRSs based on Mahajan *et al*. and Scott *et al*. Results from association tests with GDM. Abbreviations: 0 85 90 95 and 98%, PRSs including a subset of SNPs tested in at least 0 85 90 95 and 98% of the total samples of the consortium study respectively; AUC area under the curve; PRS, Polygenic Risk Score; LD, Linkage disequilibrium; P+T, Pruning and thresholding; SNP, Single Nucleotide Polymorphism; START, South Asian Birth Cohort; ROC, receiver operating characteristic; uPRS, unweighted PRSs; wPRS, weighted PRSs.

**Figure 2:**
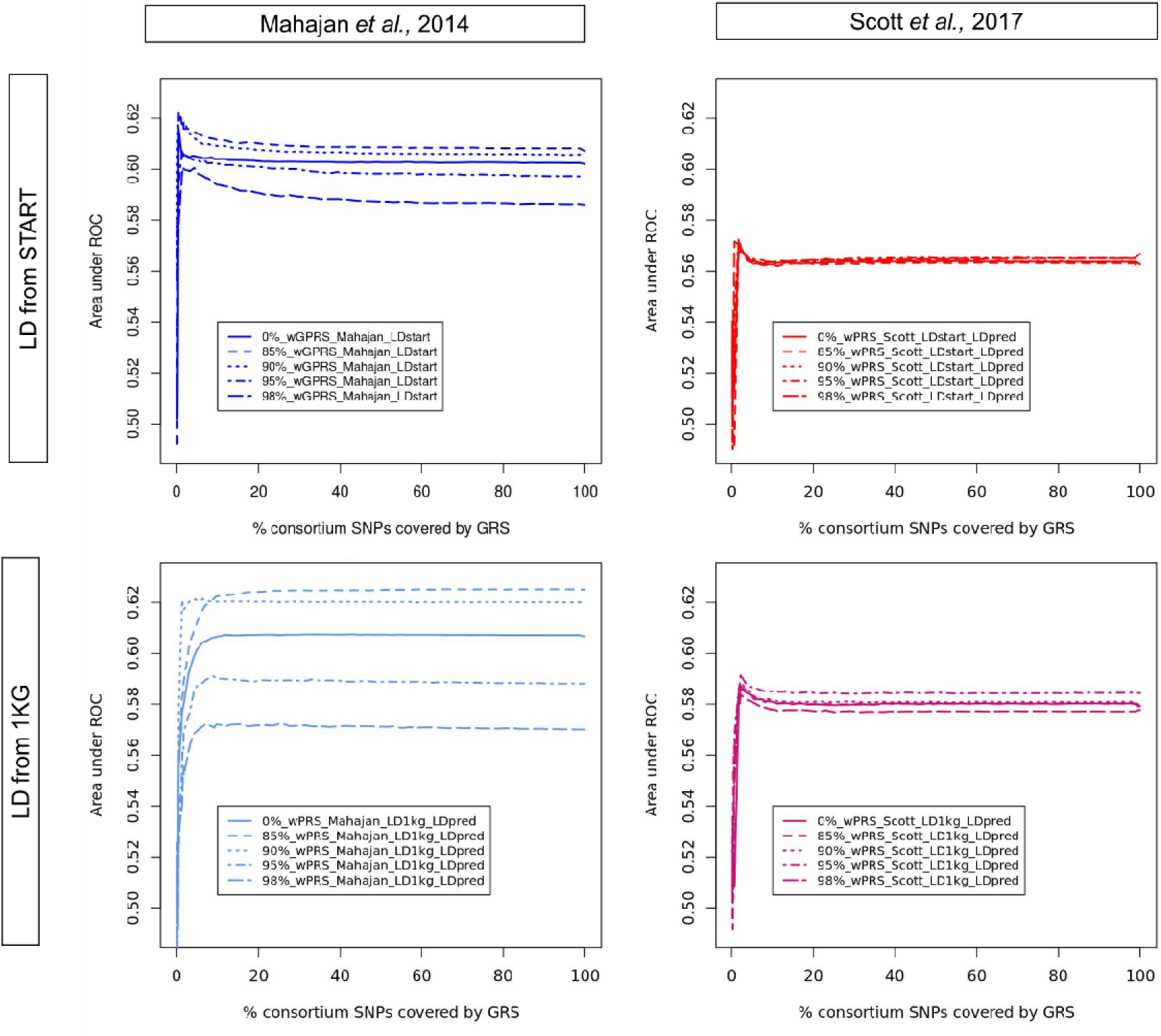
AUCs of the different LDpred PRSs based on Mahajan *et al*. and Scott *et al*. Results from association tests with GDM. Abbreviations: 0 85 90 95 and 98%, PRSs including a subset of SNPs tested in at least 0 85 90 95 and 98% of the total samples of the consortium study respectively; 1KG, 1000 Genomes; AUC area under the curve; PRS, Polygenic Risk Score; LD, Linkage disequilibrium; SNP, Single Nucleotide Polymorphism; ROC, receiver operating characteristic; wPRS, weighted PRSs.

With an AUC of 0.62, the best GraBLD PRS included 1,305,596 SNPs and was derived using weights from Mahajan *et al*. (W_Mahajan_); LD from START (LD_START_); SNPs tested in ≥ 90% of the samples in the consortium data (N_90%_). This top GraBLD PRS will hereafter be referred to as GraBLD_PRS_1__W_Mahajan__LD_START__N_90%_. The best P+T was comprised of 9,274 SNPs, showed an AUC of 0.62 and was derived using the following parameters: Weights from Mahajan *et al*. (W_Mahajan_); LD from 1KG (LD_1KG_); SNPs tested in ≥ 85% of the maximum sample size (N_85%_) and a maximum P-value of 0.016 in its reference consortium GWAMA (P_0.016_). This top P+T PRS will be referred to as PT_PRS_1__W_Mahajan__LD_1KG__N_85%min__P_0.016._ Finally, with an AUC of 0.62 as well, the best LDpred PRS included 1,290,525 SNPs and was derived using weights from Mahajan *et al*. (W_Mahajan_); LD from 1KG (LD_1KG_); SNPs tested in ≥ 85% of the samples in the consortium data (N_85%_). This top LDpred PRS will hereafter be referred to as LDpred_PRS_1__W_Mahajan__LD_1KG__N_85%_. Detailed characteristics and rankings of the best PRSs are shown in Supplementary Table 2

### Pruning and Threshold PRSs

AUCs of unweighted P+T PRSs were similar to AUCs for their weighted counterpart at very low P-value thresholds (clump P ≤ 0.004). Interestingly, the inclusion of variants with association P-values that are less significant than the usual GWAS significance threshold (5 × 10^−8^) always resulted in a considerable increase in the predictive value of the scores. Optimal AUCs were reached at clump P-values ranging from 0.016 – 0.20 depending on the source of LD or the consortium used (Figure 1, Supplementary Figure 2 and Table 2). Passed these maximal points, the difference between weighted and unweighted PRSs gradually increased, with the weighted PRSs performing better than their unweighted counterparts (Figure 1, Supplementary Figure 2).

### LDpred PRSs

Similarly to P+T PRSs, the increase in the fraction of causal SNPs (from P values of 5 × 10^−8^ to P = 0.0005 corresponding to 0.5 and 7.5 % of consortium SNPs in Mahajan *et al*. and Scott *et al*. respectively) highly improved the predictive value of the PRSs (Figure 2). The increase in the fraction causal passed this point was not associated with a significant change in the AUC of LDpred PRSs (Figure 2).

#### GraBLD vs. P+T vs. LDpred PRSs

As previously mentioned, whether the performance of a PRS derived using a given method was better than that of its different counterparts (other two methods) largely depended on the consortium data, the source of LD, the minimum % of participants, and the maximum clump P-value cutoffs used. Nevertheless, when comparing the best PRSs derided from each method, no significant difference was observed between GraBLD, LDpred and P+T (AUCs=0.62, Table 2, P _pairwise differences_ = 0.95). When comparing P+T to LDpred only, AUCs were higher and more stable in LDpred PRSs than in P+T PRS for high P-value thresholds (> 0.1) (Figure 3).

**Figure 3:**
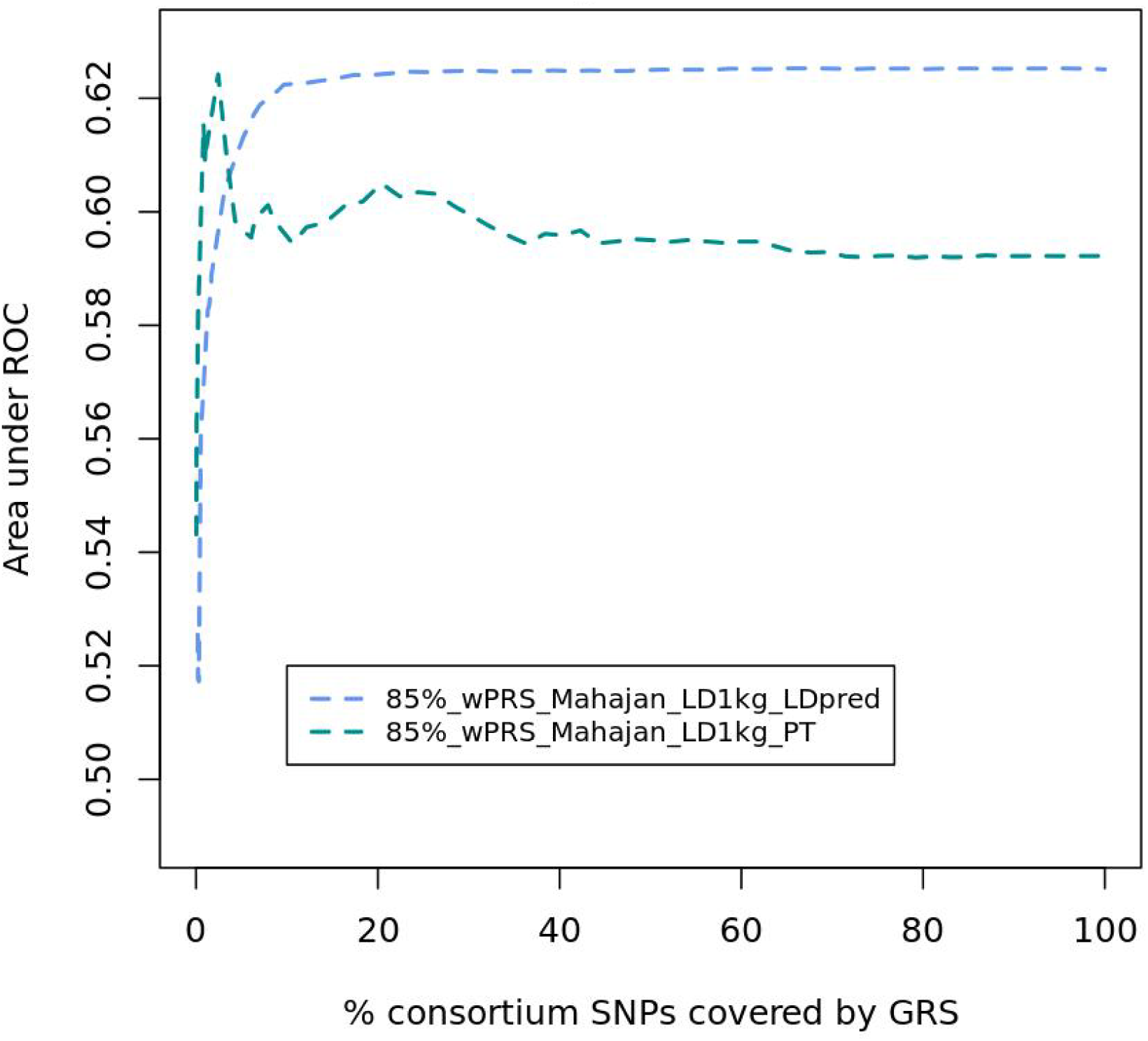
AUCs of the best Mahajan *et al*. P+T and LDpred PRSs in START. Abbreviations: 85 %, PRSs including a subset of SNPs tested in at least 85 % of the total samples of the consortium study respectively; 1KG, 1000 Genomes; AUC area under the curve; PRS, Polygenic Risk Score; LD, Linkage disequilibrium; P+T, Pruning and thresholding; START, South Asian Birth Cohort; ROC, receiver operating characteristic; wPRS, weighted PRSs

#### Mahajan et al. vs. Scott et al. PRSs

The predictive value of most Mahajan-based P+T PRSs were higher than that of their Scott-based counterparts (Figure 4). For GraBLD and LDpred PRSs, all Mahajan based PRSs had higher AUCs than Scott based PRSs (Figure 4, data not shown). Finally, the best Mahajan-based PRSs always outperformed the top Scott-based PRSs for all three methods (Table 2, Supplementary Table 2)

**Figure 4:**
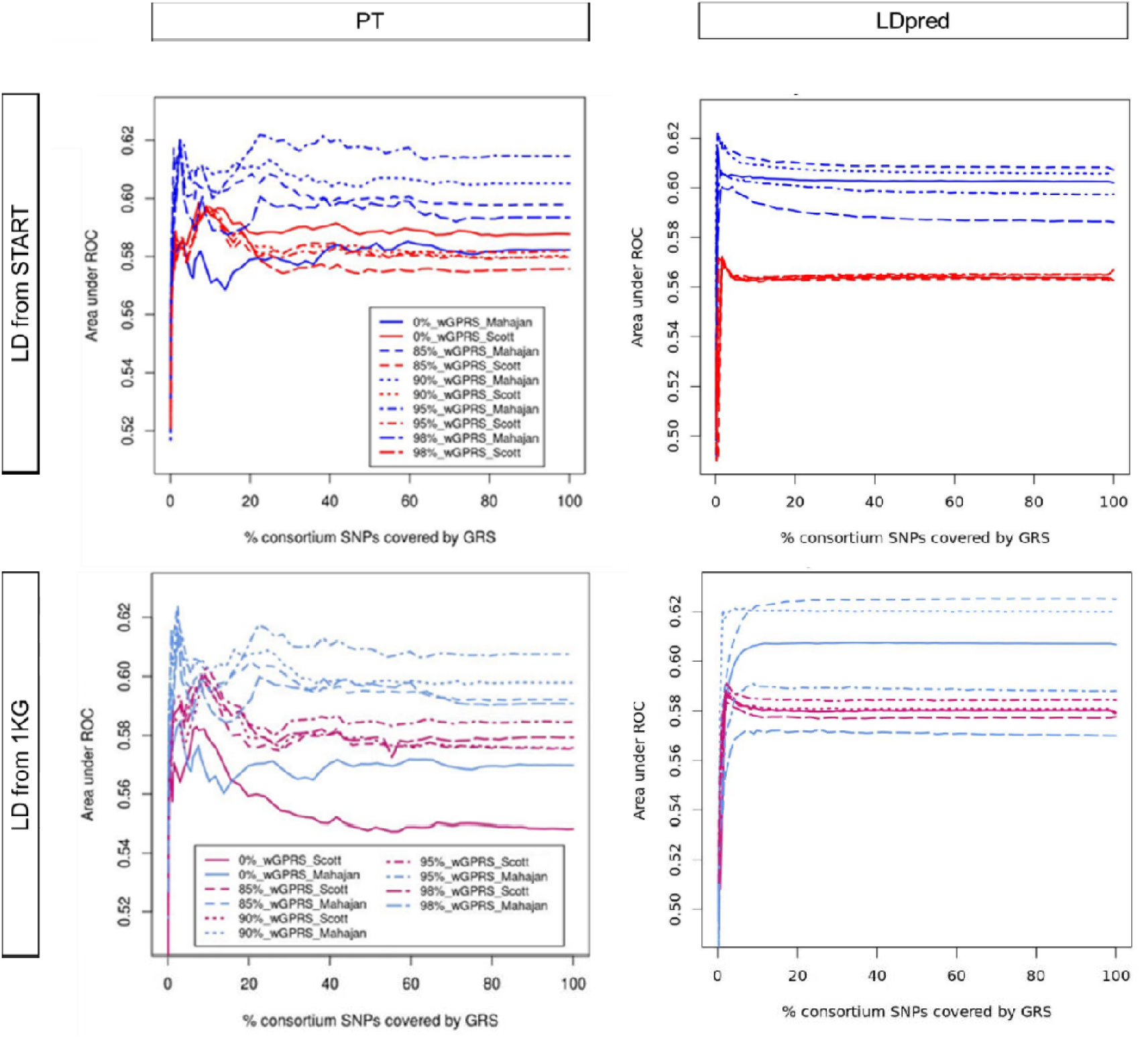
AUCs of the different weighted P+T and LDpred PRSs based on Mahajan *et al*. and Scott *et al*. Results from association tests with GDM. Abbreviations: 0 85 90 95 and 98%, PRSs including a subset of SNPs tested in at least 0 85 90 95 and 98% of the total samples of the consortium study respectively; 1KG, 1000 Genomes; AUC area under the curve; PRS, Genome-wide Polygenic Risk Score; LD, Linkage disequilibrium; P+T, Pruning and thresholding; SNP, Single Nucleotide Polymorphism; START, South Asian Birth Cohort; ROC, receiver operating characteristic; uPRS, unweighted PRSs; wPRS, weighted PRSs.

#### LD from 1000 Genomes vs. START

AUCs of the best LD_START_ PRSs were not significantly different from AUCs of their LD_1KG_ counterpart for P+T, LDpred and GraBLD PRSs (Figure 5, Table 2).

**Figure 5:**
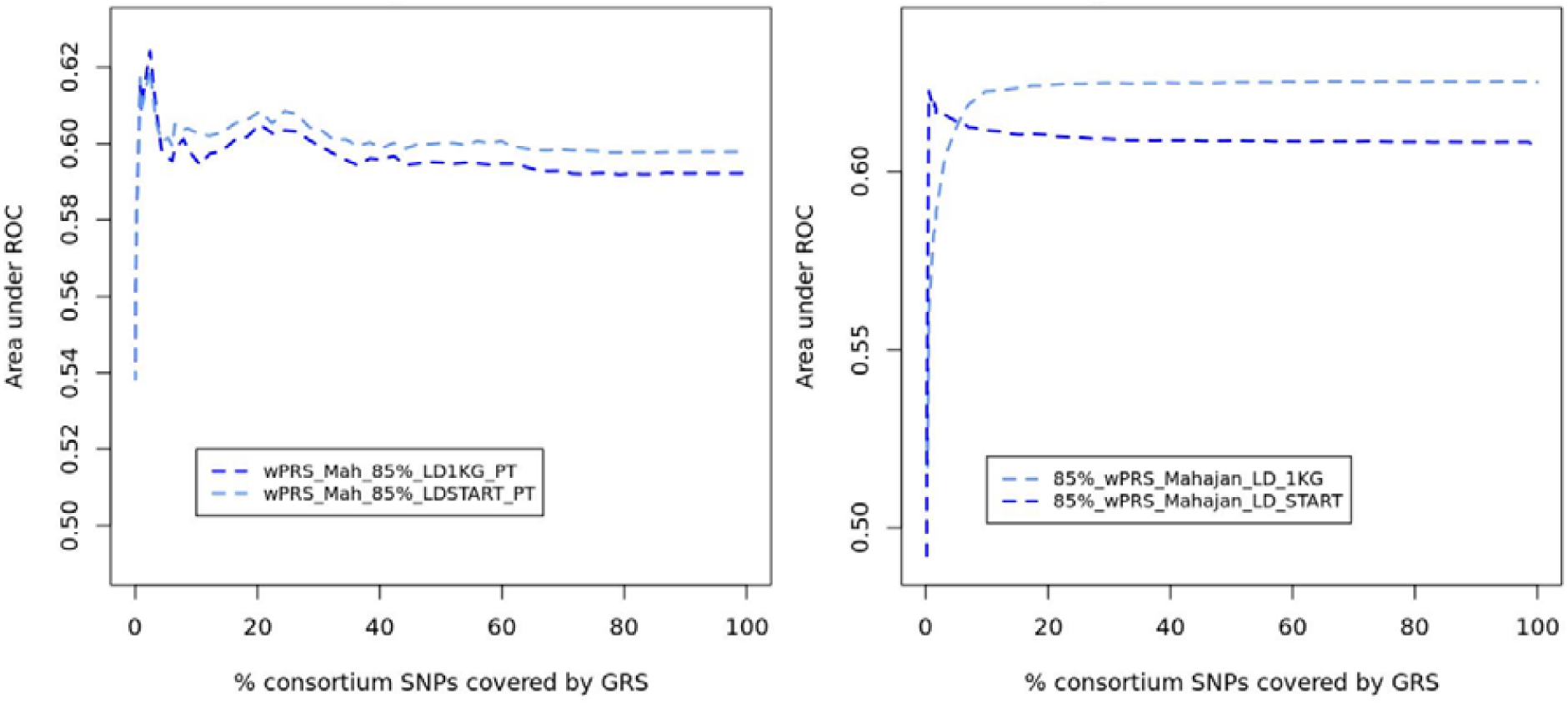
AUCs of Mahajan *et al*. N_85%_ based LD_START_ P+T and LDpred PRS and their LD_1KG_ counterparts. Abbreviations: 85 and 95%, PRSs including a subset of SNPs tested in at least 85 and 95% of the total samples of the consortium study respectively; 1KG, 1000 Genomes; AUC area under the curve; PRS, Polygenic Risk Score; LD, Linkage disequilibrium; P+T, Pruning and thresholding; START, South Asian Birth Cohort; ROC, receiver operating characteristic; wPRS, weighted PRSs.

### Association with GDM

The association results of the top PRSs (GraBLD_PRS_1__W_Mahajan__LD_START__N_90%,_ PT_PRS_1__W_Mahajan__LD_1KG__N_85%__P_0.016_ and LDpred_PRS_1__W_Mahajan__LD_1KG__N_85%_) with GDM (univariate models) are shown in Table 2 (continuous PRSs) and Table 3 (categorical PRSs). The odds of developing GDM was 2 to 2.5 fold higher in participants with the highest PRSs (top 25%) compared to the rest (75%) of the study population, depending on the type of PRS used. When analyzing participants with high and low PRSs values only, our results show that participants with the highest PRS values (top 25%) had between 3 and 3.4 fold increase in their risk of GDM compared to the participants with the lowest PRS values (bottom 25%). These results were similar in both UK Biobank South Asian and white British replication datasets (Table 3).

**Table 3:** Association results of Top PRSs (categories) with GDM in South Asian women from the START and UK Biobank studies. GraBLD PRS used: GraBLD_PRS_1__W_Mahajan__LD_START__N_90%_; P+T PRS used: PT_PRS_1__W_Mahajan__LD_1KG__N_85%min__P_0.016_; LDpred PRS used: LDpred_PRS_1__W_Mahajan__LD_1KG__N_85%_. Abbreviations: CI, confidence interval; PRS, Polygenic Risk Score; GraBLD, Gradient Boosted and LD adjusted; OR, Odds ratio; P+T, Pruning and thresholding; SAW, South Asian Women; START, South Asian Birth Cohort; WBW, White British Women.

| High PRS definition | Reference group | PRS type | START – SAW |  |  | UK Biobank – SAW |  |  | UK Biobank – WBW |  |  |
| --- | --- | --- | --- | --- | --- | --- | --- | --- | --- | --- | --- |
|  |  |  | OR | 95% CI | P value | OR | 95% CI | P value | OR | 95% CI | P value |
| Top 25% | Remaining 75% | GraBLD | 2.51 | 1.82 - 3.47 | $1.75 \times 10^{-8}$ | 2.66 | 1.51 - 4.63 | 0.0006 | 1.75 | 1.48 – 2.05 | $2.07 \times 10^{-11}$ |
| | | P+T | 2.08 | 1.51 – 2.87 | $7.44 \times 10^{-6}$ | 1.80 | 0.99 – 3.17 | 0.05 | 1.65 | 1.40 – 1.94 | $2.41 \times 10^{-9}$ |
| | | LDpred | 2.00 | 1.45 – 2.76 | $2.11 \times 10^{-5}$ | 2.61 | 1.16 – 3.60 | 0.01 | 1.72 | 1.46 – 2.02 | $7.66 \times 10^{-11}$ |
| Top 25% | Lowest 25% | GraBLD | 3.40 | 2.25 - 5.17 | $7.30 \times 10^{-9}$ | 5.30 | 2.17 - 15.88 | 0.0008 | 2.11 | 1.69 – 2.66 | $1.09 \times 10^{-10}$ |
| | | P+T | 3.09 | 2.10 – 4.74 | $1.47 \times 10^{-7}$ | 4.21 | 1.67 – 12.82 | 0.005 | 2.22 | 1.76 - 2.82 | $3.08 \times 10^{-11}$ |
| | | LDpred | 3.06 | 2.02 – 4.69 | $1.77 \times 10^{-7}$ | 3.59 | 1.53 – 9.84 | 0.006 | 2.08 | 1.66 – 2.62 | $3.09 \times 10^{-10}$ |

### Heritability

In order to better characterize the genetic architecture of GDM, heritability was estimated from 1) genome-wide genotype data (h^2^_WG_SNPs)_; 2) common variants (MAF ≥ 0.01); and 3) SNPs included in our top P+T PRS, in South Asian women from START, as well as for a subgroup of white British women from UK Biobank. The results are shown in Table 4. Due to a lack of power, standard errors for our heritability estimates were large, and most lower and upper bound values of the 95% confidence intervals were close to, or crossed the respective 0 and 1 boundaries. The proportion of the variance in GDM explained by genome-wide data (h^2^_WG_SNPs_) in South Asian women from the START study was 0.57 ± 0.88 and h^2^_WG_SNPs_ estimated in women from white British UK Biobank samples was 0.15 ± 0.22 (Table 4). Heritability attributed to common variants (MAF ≥ 1%) reached 0.5 ± 0.78 and 0.11 ± 0.14 in our START and UK Biobank samples, which explained 88% and 73% of the h^2^_WG_SNPs_ respectively. Heritability estimated from the SNPs used in our top P+T PRS explained 31.6% and 20 % of h^2^_WG_SNPs_ in START and UK Biobank respectively.

**Table 4:** Heritability estimates for GDM in women from the START and UK Biobank studies. Models are adjusted for the first 3 eigenvectors from the principal component analysis. * heritability estimates on the liability scale (disease prevalence = 0.4%) from the case-cohort study. Abbreviations: MAC, Minor allele count; MAF, Minor allele frequency; NA, Not applicable; P+T, Pruning and thresholding; SAW, South Asian women; SE, Standard Error; SNP, Single Nucleotide Polymorphism; START, South Asian Birth Cohort; WBW, White British Women.

|  |  | START - SAW |  | UK Biobank - WBW * |  |
| --- | --- | --- | --- | --- | --- |
|  | Allele count /<br>MAF cutoff | h<sup>2</sup> | SE | h<sup>2</sup> | SE |
| All SNPs in the study | MAC 10 | 0.57 | 0.43 | 0.15 | 0.11 |
|  | MAF 0.01 | 0.50 | 0.40 | 0.11 | 0.07 |
| SNPs in top P+T PRS | NA | 0.18 | 0.14 | 0.03 | 0.03 |

## DISCUSSION

In this study, we built several thousands of GDM PRS using genome-wide genotypes, large consortium data and 3 different methods. Our best PRS was built using the LDpred method, with weights extracted from Mahajan *et al*. and LD calculated using 1KG genotypes. This PRS was significantly associated with GDM in South Asian women from the START study, an observation that was successfully replicated in South Asian and white British women from UK Biobank. Participants with the highest PRS values had an increased risk of GDM when compared to the other groups.

We observed a considerable difference in the proportion of participants with GDM between South Asian women from the START study (36.2 %), South Asian women from UK Biobank (2.2%), and white British women from UK Biobank (0.43%). This disparity is likely due to major differences in the study design, recruitment strategies and definition of GDM between the two studies involved. For instance, the definition of GDM status in START was based on glucose levels measurements performed during pregnancy in response to an oral glucose challenge, and South Asian-specific diagnosis cutoffs were used. On the other hand, GDM status was retrospectively self-reported by UK Biobank participants, which most likely resulted in an increased number of misclassifications and a reduced number of reported GDM cases. In an effort to refine the phenotype in UK Biobank, our control group was restricted to women without GDM who also went through at least one live birth. Nevertheless, the lack of sensitivity and specificity of the GDM phenotype likely remains an issue in UK Biobank.

Summary statistics from two large T2D GWAMAs were used to build our PRSs. One of the major advantages in using data from Mahajan *et al*. was that ∼20% of its participants in their publically available data originated from the South Asian sub-continent. The study also had a large maximum number of cases and controls, but many of the SNPs included in the meta-analysis were tested in a much smaller sample (Supplementary figure 3, Supplementary Table 1). On the other hand, no South Asian participants were included in the GWAMA performed by Scott *et al*. but the average number of samples tested for each SNP was larger than in Mahajan *et al*. Our results show that Mahajan-based PRSs consistently outperformed their Scott-based counterparts in spite of a lower genome coverage and smaller average number of participants per SNP. This highlights the importance of using consortium data of the same ethnic group than the study at hand whenever possible. However, since Mahajan *et al*.’s summary statistics were derived from a blend of participants of different ethnicities, our top PRS could likely be improved if built based on summary statistics derived from an equally powered GWAMA performed in South Asians only.

Several reports suggest that T2D and GDM share a common genetic background. In the absence of publicly available data of large GDM GWASs, summary statistics from a T2D consortium were used to derive our scores. Our results show that a T2D PRSs can be highly predictive of GDM in South Asian and European-origin women, hence confirming the hypothesis of a common genetic background between these two diseases. However, the effect size of the genetic variants could be different between the two conditions, and some loci could be specific to each disease. Although these differences should not affect our models comparisons, we expect that the predictive value of GDM PRSs will be further improved if built using weights from a large GDM GWAS or GWAMA.

A significant conclusion derived from this study is that, whatever the consortium or the method used, restricting the list of SNPs to GWAS significant variants (P value ≤ 5×10^−8^) drastically reduces the predictive value of the PRSs. Unfortunately, many studies still rely on this threshold to select their loci of interest to derive their risk scores. We recommend the use of higher P-value thresholds (> 0.01 in our case) whenever possible in order to increase the predictive value of the PRSs.

Based on our results, weighted PRSs perform similarly, or better than their unweighted counterparts in general. For P+T PRS, the predictive value of the unweighted PRSs are especially lower than for weighted PRS at high P-value cutoffs. Hence, we recommend the use weighted PRSs whenever possible. If unweighted PRSs are used, P-value cutoffs should be set between 0.01 and 0.1.

When comparing the best PRSs, our results suggest that the GraBLD, P+T and LDpred methods perform equally well in terms of disease prediction as measured by the AUC. Nevertheless, the identification of the optimal P+T, and LDpred PRSs required the test of several thousand predictors (n = 2,560 and 1280 respectively), when a similar result was achieved by testing 40 GraBLD models only. While this may lead one to slightly favor the use of LDpred or GraBLD for the building of PRSs, the P+T remains the method of choice in our opinion, given the fact that it required less SNPs and was easier to implement using the PLINK software.

T2D’s SNP-based heritability has recently been estimated at 0.54 [95%CI: 0.47 – 0.61] ^51^. Our results from START suggest that the heritability of GDM is similarly high (h^2^ _WG_SNPs_ in START=0.58, SE=0.43) although the standard error is large. Heritability estimates were considerably smaller in European participants in UK Biobank. This could be explained by the difference between the disease prevalence in START and UK Biobank as previously discussed, and potentially, by differences in the environments and lifestyles of the participants included in the two studies. Another potentially interesting observation derived from our heritability results is that a large proportion of the genetic variance of GDM (between 73.3% and 87.7% depending on the ethnicity) can be explained by common SNPs. Furthermore, our results suggest that a relatively large proportion of the genetic variance (between 20% and 31.6%) is captured by an even smaller fraction of SNPs (∼2.5%, N=9,274) that are included in our top P+T PRS. However, given the small number of cases in our studies, our GREML tests are likely underpowered, resulting in very large standard errors. Hence, our heritability results should be interpreted with caution, and larger studies are needed in order estimate the heritability of GDM with higher accuracy. An interesting future analysis would be to further partition these heritability estimates into maternal and fetal genetic effects. Indeed, since maternal glucose cut-offs used to define GDM also rely on the fetal phenotypes (size and adiposity related measures), we strongly suspect that the fetus’s genome will also contribute to the mother’s risk of developing the disease. Unfortunately, this analysis could not be performed in our study because of our small sample size.

In conclusion, our results show that the predictive value of polygenic risk scores in South Asian women can be greatly improved by combining genome-wide genotyping data and by extracting summary statistics from large multi-ethnic genome-wide meta-analysis.

## METHODS

### Study design and participants

#### The South Asian Birth Cohort (START) study

START is a prospective cohort designed to evaluate the environmental and genetic determinants of cardiometabolic traits of South Asian pregnant women and their offspring living in Ontario, Canada. The rationale and study design are described elsewhere ^52^. In brief, 1,012 South Asian (people who originate from the Indian subcontinent) pregnant women, between the ages of 18 and 40 years old, were recruited during their second trimester of pregnancy from the Peel Region (Ontario, Canada) through physician referrals between July 11, 2011 and Nov. 10, 2015. All START participants signed an informed consent including genetic consent, and the study was approved by local ethics committees (Hamilton Integrated Research Ethics Bard, William Osler Health System, and Trillium Health Partners). A detailed description of the maternal measurements has been published previously ^53^.

#### UK Biobank

The UK Biobank is a large population-based study which includes over 500,000 participants living in the United Kingdom ^50^. Men and Women aged 40 – 69 years were recruited between 2006 and 2010 and extensive phenotypic and genotypic data about the participants was collected, including ethnicity and a question regarding past history of GDM. Details of this study are available online (https://www.ukbiobank.ac.uk) ^50^. Data from UK Biobank were used in order to validate the results from the START study.

### Derived variables

#### START Study

GDM status was determined using the South Asian specific cutoffs as defined in the Born in Bradford study (fasting glucose level of 5.2 mmol/L or higher, or a 2-hour post load level of 7.2 mmol/L or higher) ^54^. Self-reported GDM status was used if these measures were unavailable. Participants with a history of T2D prior to pregnancy were excluded. Using these criteria, 832 START participants with known GDM status (301 cases and 531 controls) and available genotypes were included in the analysis. The South Asian ethnicity/ancestry of participants was validated using genetic data.

#### UK Biobank

Participants in the UK Biobank completed questionnaires at several time points (questionnaire of initial assessment visit, 2006-2010; questionnaire of first repeat assessment visit, 2012-2013; questionnaire of imaging visit, 2014 onwards). For the purpose of our study, GDM cases were defined as women who reported having had diabetes during pregnancy only, collected at any time point using questionnaires. The control group was comprised of women who: 1) had at least one child (self-reported, live births only) 2) had never been diagnosed with diabetes or GDM in all assessments. The ethnicity/ancestry of participants was validated using genetic data.

### DNA extraction, genotyping, imputation, filtering and SNP extraction

#### START

DNA was extracted and genotyped from a total of 867 samples (START mothers) using the Illumina Human CoreExome-24 and Infinium CoreExome-24 arrays (Illumina, San-Digeo, CA, USA). Data was cleaned using standard quality control (QC) procedures ^55^ and 837 women samples passed the QC. Genotypes were subsequently phased using SHAPEIT v2.12 ^56^, and imputed with the IMPUTE v2.3.2 software ^57^, using the 1000 Genomes (phase 3) data as a reference panel ^49^. Variants with an info score ≥ 0.7 were kept for analysis. Addition data manipulation and SNP selection criteria for the building of the PRSs are detailed in Supplementary Information and Supplementary Figure 1.

#### UK Biobank

A total of ∼500,000 participants from the UK biobank were genotyped using the UK BiLEVE or UK Biobank Affymetrix Axiom arrays. Detailed QC, phasing and imputation procedures have previously been described ^58^. As a result, 3,169 and 220,703 unrelated South Asian and white British women respectively passed QC. Among these, 2,386 and 184,869 participants had available GDM status respectively, and were used to replicate our results from the START study. Genotypes for > 98% of SNPs included in our top START GDM PRSs were available (info score ≥ 0.6) and were extracted for the replication.

### Consortium data

Summary statistics (P-value, effect size) from the following two DIAGRAM sources were used in order to build the PRSs:

#### 1) Mahajan et al., Nature Genetics, 2014

This trans-ethnic GWAMA included up to 12,171 T2D cases and 56,862 controls of European ancestry; 6,952 cases and 11,865 controls of East Asian ancestry; 5,561 cases and 14,458 controls South Asian ancestry and 1,804 cases and 779 controls of Mexican and Mexican American ancestry ^5^. This study was selected for its multi-ethnic composition and its inclusion of South Asian participants.

#### 2) Scott et al.,Diabetes, 2017

This study combines data from 18 GWASs, for a total of 26,676 T2D cases and 132,532 controls of European ancestry ^6^. This study was selected for its large sample size and its relatively homogenous samples (100% white Caucasians). Summary statistics of these studies were downloaded from DIAGRAM’s main website (http://www.diagram-consortium.org).

### Templates for LD calculation

LD calculations used to build the PRSs were derived from the following two genotyping datasets: 1) START study (LD source hereafter referred to as LD_START_); and 2) the 1000 Genomes consortium (LD source hereafter referred to as LD_1KG_) phase 3. Genotypes of 1000 Genomes participants were downloaded from the project’s data portal (http://www.internationalgenome.org), and a subset of participants was created in order to match the proportion of the ethnicities represented in each consortium study.

### Pruning and thresholding PRSs

Both weighted and unweighted PRSs were built using GNU Parallel ^59^ and PLINK v1.9 (https://www.cog-genomics.org/plink2). 64 different clump P-value cutoffs ranging from 5 × 10^−8^ to 1 were tested in order to identify the optimal index variant’s significance threshold. All other parameters were set to default. A diagram of the different P+T PRSs built is show in Supplementary Figure 2.

### LDpred

LDpred PRSs were derived using the LDpred software v0.9.9 (https://github.com/bvilhjal/ldpred) ^47^. The fractions of causal variants assumed a prior were similar to the P value thresholds used for the P+T PRSs. Since the number of SNPs was different between the PRSs, The LD radius was adjusted accordingly in each model using the recommended formula (N SNP/3000). All other parameters were kept on their default setting. A diagram of the different P+T PRSs built is shown in Supplementary Figure 2.

### GraBLD

GraBLD PRSs were built using several functions available in the GraBLD R package (https://github.com/GMELab/GraBLD) ^48^. Data of all the women participating in the START study were used for the calibration. All parameters were set to default.

### Association analysis

The association of each PRS with GDM was assessed using a univariate logistic regression model, and areas under the receiver-operating characteristic (ROC) curves (AUCs, c-statistics) were compared in order to determine the PRS with the highest predictive value of GDM. Continuous PRSs were also divided into quartiles in order to compare the participants with highest PRS values to the other groups. Statistical significance of the difference between the predictive values of two PRSs was tested using the DeLong’s test for two correlated ROC curves. Analyses were performed using GNU Parallel ^59^ and R v3.3 ^60^.

### Heritability

The proportion of the variance of GDM explained by: 1) genome-wide genotypes (minor allele count ≥10); 2) common variants (MAF ≥ 0.01); 3) SNPs included in our top P+T PRS; was estimated in unrelated participants (identity by state (IBS) < 0.05) by using the GCTA software ^61-64^. Single component GREML models were tested. Because of the large number of UK Biobank white British participants, heritability of GDM was estimated in subgroup of participants selected for inclusion in a case-cohort study (N = 623 cases of GDM and 9,037 controls). Hence, reported values for this study were adjusted for a disease prevalence of 0.4%, as estimated in all of the white British women with GDM data in UK Biobank (Table 1). Heritability was not estimated in South Asian women from UK biobank given the small number of GDM cases in this group. All models were adjusted for the first 3 PCA axes.

## ACKNOWLEDGEMENTS

The authors thank the UK Biobank for making their data available. The South Asian Birth Cohort (START) study data were collected as part of a bilateral program funded by the Indian Council of Medical Research / Canadian Institutes of Health Research (Project grant number: 298104, Foundation Scheme grant number: FDN-143255, Study grant numbers: INC 109205). This study was also supported by Heart and Stroke Foundation of Canada (grant NA7283. Sonia Anand is supported by a Tier 1 Canada Research Chair in Ethnic Diversity and Cardiovascular Disease, and a Heart and Stroke Foundation/Michael G. DeGroote Chair in Population Health Research at McMaster University. Guillaume Paré is supported by a Canada Research Chair.

## AUTHOR CONTRIBUTIONS

A.L performed all the analysis, wrote the main manuscript and prepared all figures and tables. S.M. provided guidance for the GRABLD analysis. D.D is the study coordinator for the START birth cohort, and she provided comments on the manuscript. M.G is a START co-principal investigator and he provided comments on the manuscript. G.P provided guidance for all analyses and reviewed the manuscript. S.A is a START co-principal investigator. She oversaw the development, analysis, and writing of this manuscript.

## ABBREVIATIONS

1KG: 1000 Genomes
AUC: Area Under the Curve
BMI: Body Mass Index
CI: Confidence Interval
DIAGRAM: DIAbetes Genetics Replication and Meta-analysis
GDM: Gestational Diabetes
PRS: Polygenic Risk Score
GraBLD: Gradient Boosted and LD adjusted
GRM: Genomic Relationship Matrix
GREML: Genomic relatedness matrix residual maximum likelihood
GRS: Genetic Risk Score
GWAMA: Genome-Wide Association Meta-Analysis
GWAS: Genome-Wide Association Study
LD: Linkage Disequilibrium
LDMS: LD and MAF stratified
MAF: Minor Allele frequency
MAGIC: Meta-Analyses of Glucose and Insulin-related traits Consortium
MS: MAF stratified
OR: Odds Ratio
PCA: Principal Component analysis
P+T: Pruning and Thresholding
ROC: Receiver Operating Characteristic
SC: Single component
SE: Standard Error
SNP: Single Nucleotide polymorphism
START: South Asian Birth Cohort
T2D: Type 2 Diabetes

