## Supplementary Data for "Genome-Wide Polygenic Risk Scores and Prediction of Gestational Diabetes in South Asian Women"

### Supplementary Information

**Data manipulation and SNP selection:**

**START Study:** Coordinates of SNPs from Mahajan *et al.* were converted from the human genome assembly hg18 (Mar. 2006) to hg19 (Feb. 2009) using spell out UCSC's LifOver tool (UCSC) [56]. Consortium variants that met the following criteria were kept if there was the presence of: 1) valid hg19 genomic coordinates; 2) a MAF ≥0.01 in START; 3) available genotypes in START and 1000 Genomes datasets (SNPs were matched based on their hg19 chromosome, positions and alleles); and if the had been 4) tested in South Asian samples (for the GRSs based on Mahajan *et al.* only).

Additional GRSs were created and for which only SNPs tested in ≥ than 85, 90, 95% and 95% of samples (maximum sample size in the study) were kept. A detailed diagram of SNP selection process for each GRS is shown in Supplementary Figure 1.

| | Mahajan *et al.,* 2014 | | | Scott *et al.,* 2017 | | |
|---|---|---|---|---|---|---|
| | Minimum N samples tested per SNP | N SNPs remaining after filtering | % SNP loss from ref | Minimum N samples tested per SNP | N SNPs remaining after filtering | % SNP loss from ref |
| Keep All SNPs (ref) | 25 | 2,324,032 | 0 | 4,731 | 6,813,331 | 0 |
| Remove SNPs tested in <85% of samples | 936,81 | 1,602,182 | 31.1 | 134,452 | 6,617,007 | 2.9 |
| Remove SNPs tested in <90% of samples | 99,192 | 1,305,771 | 43.8 | 142,362 | 6,483,925 | 4.8 |
| Remove SNPs tested in <95% of samples | 104,703 | 346,290 | 85.1 | 150,275 | 6,146,154 | 9.8 |
| Remove SNPs tested in <98% of samples | 108,010 | 223,912 | 90.4 | 155,174 | 5,301,848 | 22.2 |

**Supplementary Table 1:Minimum sample size, number of SNPs to be tested in the GPRSs, and percentage of SNP loss for the selected sample size thresholds in data derived from Mahajan *et al.* and Scott *et al.*** Abbreviations: ref, reference; SNP, Single nucleotide polymorphism.

| Method | Consortium | LD source | Min % participants | P value threshold | N Variants included in the PRS | % SNPs covered by PRS | Rank (within method) * | Rank (all PRSs) * |
|---|---|---|---|---|---|---|---|---|
| PT | Mahajan et al, 2014 | 1KG | 85% | 0.016 | 9,274 | 2.42 | 1 | 106 |
| | | START | 95% | 0.200 | 35,274 | 22.34 | 3 | 307 |
| | Scott et al, 2017 | 1KG | 95% | 0.071 | 73,130 | 9.88 | 211 | 1077 |
| | | START | 0% | 0.048 | 67,693 | 7.07 | 278 | 1237 |
| GraBLD | Mahajan et al, 2014 | 1KG | 90% | NA | 1,305,596 | NA | 65 | 107 |
| | | START | 90% | NA | 1,305,596 | NA | 1 | 18 |
| | Scott et al, 2017 | 1KG | 98% | NA | 5,302,459 | NA | 961 | 1894 |
| | | START | 98% | NA | 5,302,459 | NA | 641 | 1458 |
| LDpred | Mahajan et al, 2014 | 1KG | 85% | 0.94 | 1290525 | 94.64 | 1 | 1 |
| | | START | 90% | 1 | 1305596 | 100 | 47 | 241 |
| | Scott et al, 2017 | 1KG | 95% | 0.01 | 112875 | 2.25 | 426 | 1800 |
| | | START | 95% | 0.007 | 88472 | 1.70 | 834 | 3199 |

**Supplementary Table 2: characteristic of the best PRS for each method, consortium data and LD source used**. *weighted PRS considered in the ranking. Abbreviations: 1KG, 1000Genomes; AUC area under the curve; PRS, Polygenic Risk Score; GraBLD, Gradient Boosted and LD adjusted; LD, Linkage Disequilibrium ;Min, minimum; NA, Non applicable; P+T, pruning and thresholding; SNP, Single Nucleotide Polymorphism;

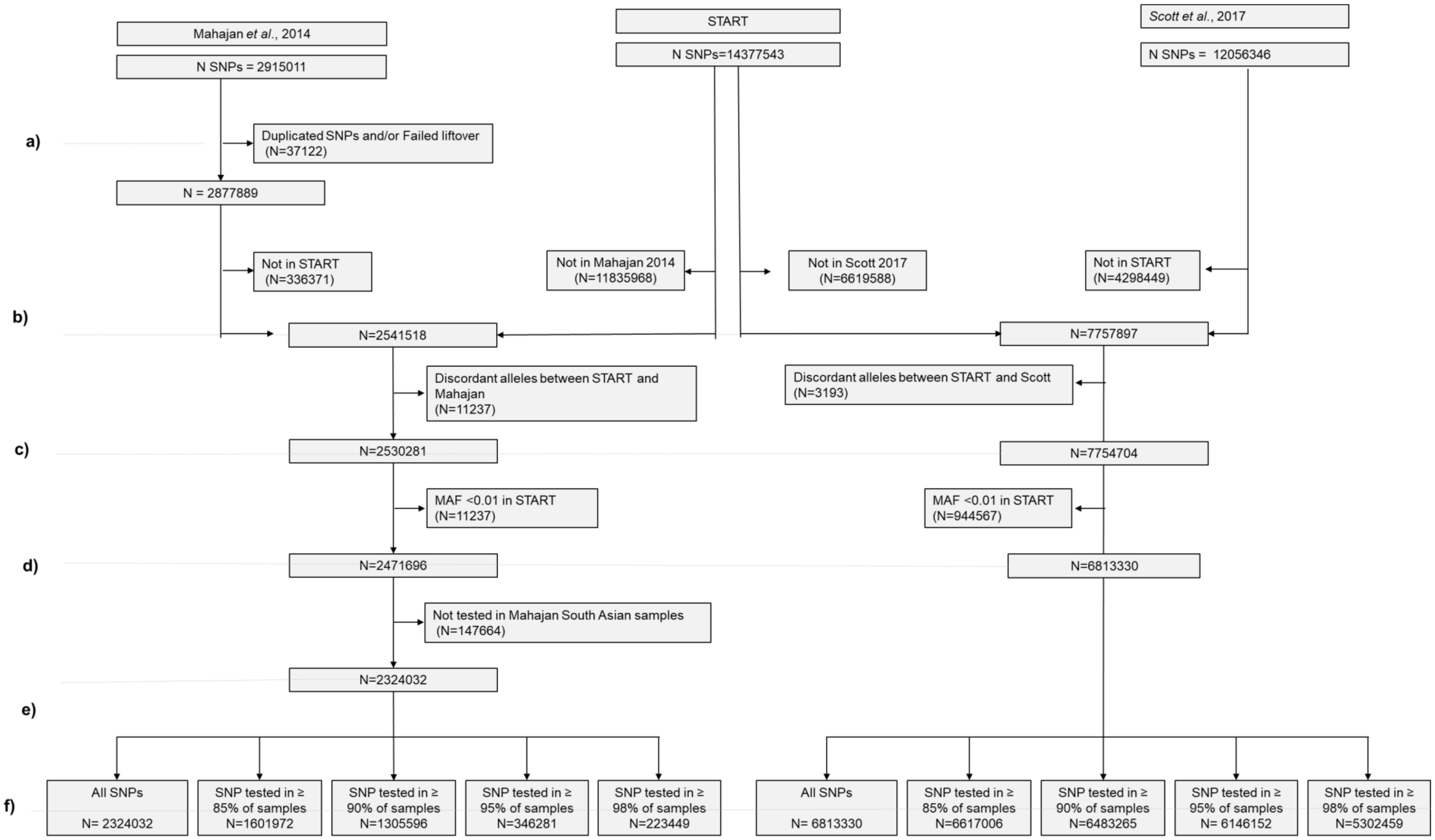


**Supplementary Figure 1: Diagram of SNP filtering steps prior to building the GPRSs in START.** Abbreviations: MAF, minor allele frequency; SNP, Single nucleotide polymorphism; START, South Asian birth cohort.

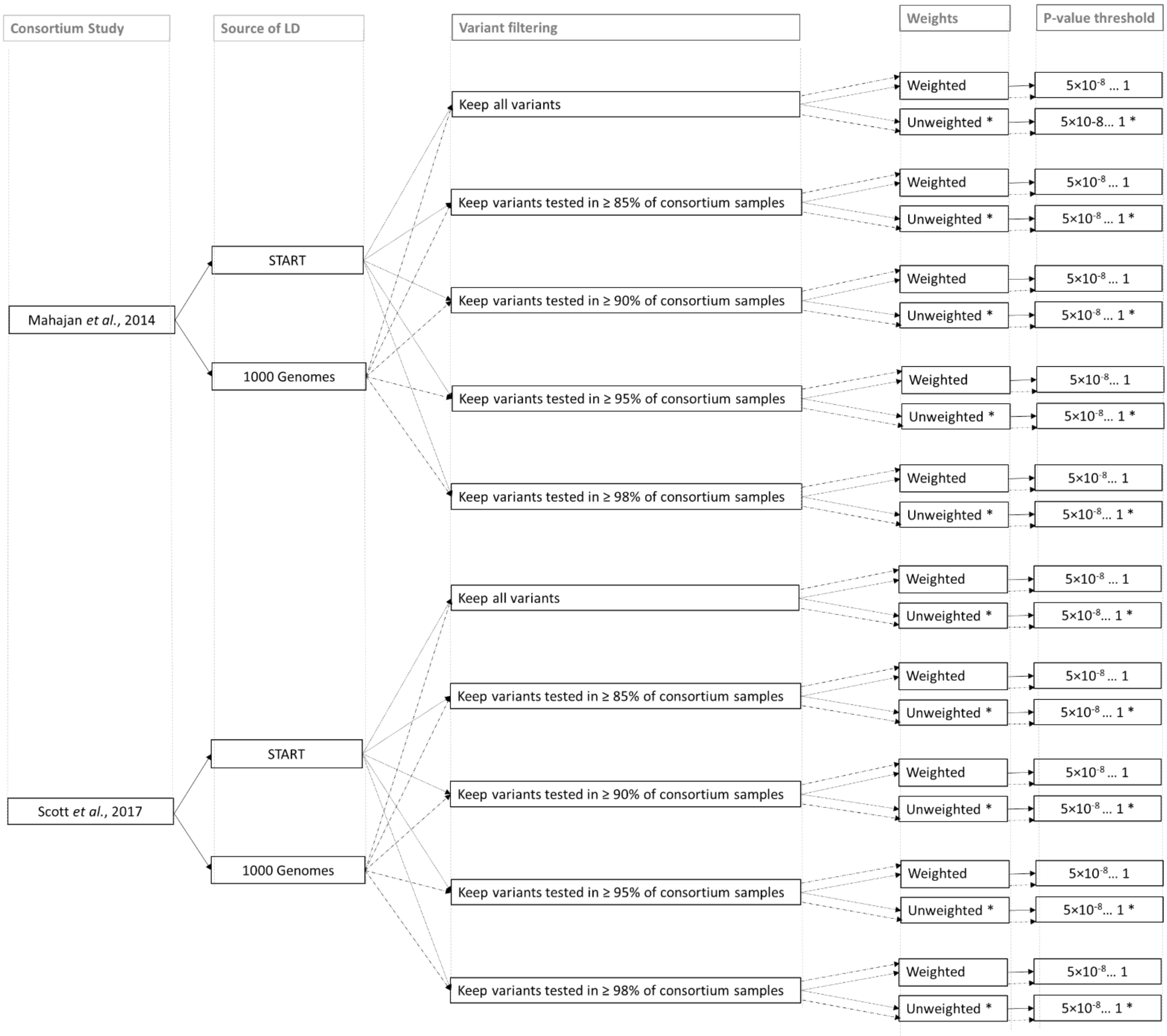


**Supplementary Figure 2: Diagram of the different T2D P+T and LDpred PRSs derived for South Asian women from the START study.** *, applies to P+T PRSs only. Abbreviations: START, South Asian birth cohort.

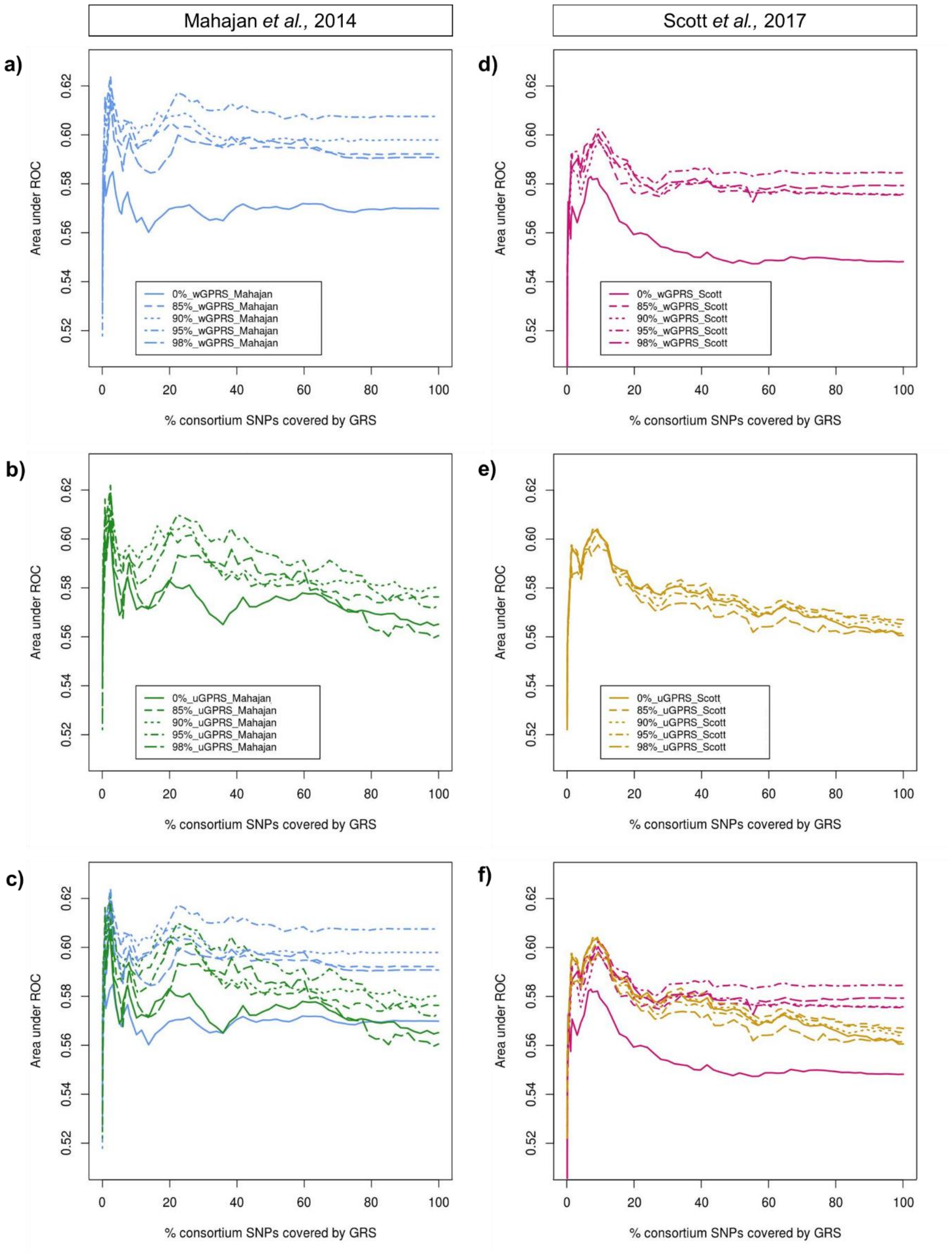


**Figure 2: AUCs of the different weighted and unweighted $LD_{1KG}$ P+T PRSs based on Mahajan *et al.* and Scott *et al.*** Results from association tests with GDM. Abbreviations: 0 85 90 95 and 98%, PRS including a subset of SNPs tested in at least 0 85 90 95 and 98% of the total samples of the consortium study respectively; 1KG, 1000 Genomes; AUC area under the curve; PRS, Genome-wide Polygenic Risk Score; LD, Linkage disequilibrium; P+T, Pruning and thresholding; SNP, Single Nucleotide Polymorphism; ROC, receiver operating characteristic; uPRS, unweighted PRS; wPRS, weighted PRS.

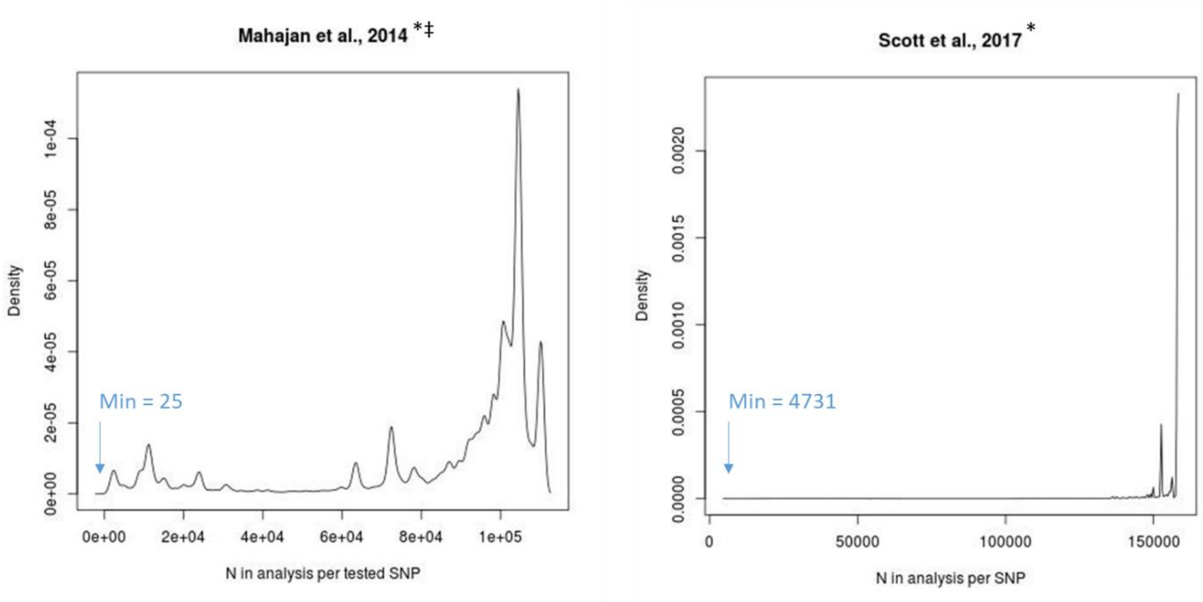


**Supplementary Figure 4: Density plot of the number of participants tested for association with T2D per SNP in Mahajan *et al.* and Scott *et al.*.**[*] SNPs common between the START and the consortium data. ‡, SNPs tested in South Asians in Consortium. Abbreviations: START, South Asian birth cohort; T2D, type 2 diabetes.
